# Agricultural fungicides inadvertently influence the fitness of Colorado potato beetles, *Leptinotarsa decemlineata*, and their susceptibility to insecticides

**DOI:** 10.1101/286377

**Authors:** Justin Clements, Sean Schoville, Anna Clements, Dries Amezian, Tabatha Davis, Benjamin Sanchez-Sedillo, Christopher Bradfield, Anders S. Huseth, Russell L. Groves

**Author notes:** To whom correspondence should be addressed: Russell L. Groves, Department of Entomology, University of Wisconsin, 1630 Linden Drive, Madison, WI 53706.

## Abstract

The Colorado potato beetle (CPB), *Leptinotarsa decemlineata* (Say), is an agricultural pest of solanaceous crops which has developed insecticide resistance at an alarming rate. Up to this point, little consideration has been given to unintended, or inadvertent effects that non-insecticide xenobiotics may have on insecticide susceptibility in *L. decemlineata*. Fungicides, such as chlorothalonil and boscalid, are often used to control fungal pathogens in potato fields and are applied at regular intervals when *L. decemlineata* populations are present in the crop. In order to determine whether fungicide use may be associated with elevated levels of insecticide resistance in *L. decemlineata*, we examined phenotypic responses in *L. decemlineata* to the fungicides chlorothalonil and boscalid. Using enzymatic and transcript abundance investigations we also examined modes of molecular detoxification in response to both insecticide (imidacloprid) and fungicide (boscalid and chlorothalonil) application to more specifically determine if fungicides and insecticides induce similar metabolic detoxification mechanisms. Both chlorothalonil and boscalid exposure induced a phenotypic, enzymatic and transcript response in *L. decemlineata* which correlates with known mechanisms of insecticide resistance.

**Key Messages:**

- Prior-exposure to a fungicidal application changes the phenotypic response of *Leptinotarsa decemlineata* to the insecticide imidacloprid
- Both a fungicide and insecticide application activates similar molecular mechanisms of detoxification in *Leptinotarsa decemlineata*
- Fungicidal xenobiotics may contribute to insecticide resistance in *Leptinotarsa decemlineata*

**Author Contribution Statement:** JC, SS, CB, AH, and RG conceived and designed research. JC, AC, DA, TD, and BS conducted experiments. JC and AC analyzed data. JC and RG wrote the manuscript. All authors read and approved the manuscript.

## Introduction

The Colorado potato beetle, *Leptinotarsa decemlineata* (Say), is a major agricultural pest which causes significant crop loss and direct damage to commercial potatoes (*Solanum tuberosum*), tomatoes (*Solanum lypcopersicum*), eggplants (*Solanum melongena*), and peppers (*Solanum annuum*). The impact of *L. decemlineata* direct damage to crops is far ranging and populations of these beetles have become a significant pest throughout Asia, Europe, and North America ^1,2^. The history of insecticidal inputs for control of *L. decemlineata* is a story retold in many potato production regions of the country, where many classes of insecticides have been effective for short periods of time before the beetles become resistant. Recent estimates suggest that select populations of beetles have now become resistant to more than 56 insecticidal chemistries ^3^, and this resistance has been observed in many potato production regions of the United States of America (US) ^2^. However, the western potato production areas of the US (CO, ID, OR, and WA) are the exception, where susceptibility to several insecticide mode of action (MoA) classes remains in field populations of *L. decemlineata*. Over the past two decades (1995 to present), the registration and use of neonicotinoid insecticides (Insecticide Resistance Action Committee’s, (IRAC), MoA Group 4A compounds), including the active ingredients imidacloprid, thiamethoxam, clothianidin, and dinotefuran (all competitive modulators of the nicotinic acetylcholine receptor), has become widespread in potato ^2,4^. Since the initial introduction of neonicotinoids in the mid-1990s, *L. decemlineata* populations have steadily developed resistance to these insecticides. Nevertheless, neonicotinoids remain one of the principal insecticidal tools for control of *L. decemlineata* in potatoes throughout much of the US production regions ^5,6^. Previous studies have documented the mechanisms by which this insect rapidly develops resistance ^7–11^, and multiple biological mechanisms of resistance have been classified among tested populations. However, no study to date has fully examined all of the potential alternative drivers which may contribute to this rapid development of resistance.

While insecticides are often applied to field populations of *L. decemlineata* frequently and successively, other chemical inputs may play a role in the development of insecticide resistance. One notable factor may be cross-resistance between insecticides and fungicides that facilitates rapid evolutionary change. Cross-resistance refers to an insect’s development of tolerance or reduced sensitivity to a usually toxic, insecticidal substance as a result of exposure to a different, sub-lethal substance which may be less toxic, or non-lethal. For example, cross-resistance may be the product of the regulation of nonspecific enzymes which attack functional groups rather than the specific molecules ^12^. While *L. decemlineata* cross-resistance has previously been examined between multiple insecticides ^5,7^, no studies to date have explored the potential for cross-resistance between insecticides and fungicides, which are frequently co-applied to potato crops. If such cross-resistance does occur between select fungicides and insecticides used in potato crop culture, the genes activated could lead to more prevalent, or hastened insecticide resistance.

Patterson et al. found that exposure to the fungicide phosphite (Phostrol®, Nufarm Americas Inc., Alsip, IL) had a negative effect on *L. decemlineata* larval fitness and survival when larvae were fed treated leaves ^13^. In particular, the authors reported that the phosphite-treated potato extended larval development times and increased immature mortality. These results complement our working hypothesis and suggest that select fungicides may represent a selection factor, substantiating an inadvertent link between *L. decemlineata* insecticide susceptibility and fungicide use patterns that favor selection for resistant individuals via direct selection pressure ^13^. Obear et al. further demonstrated in the soil-dwelling grub species, *Popillia japonica*, that genes known to metabolize insecticides are activated in the presence of fungicides, such as glutathione S-transferase (GST), an enzyme that can catalyze the conjugation of glutathione to a xenobiotic compound rendering it more easily excreted ^14^. This result also demonstrates the potential for both fungicides and insecticides to activate similar genetic pathways of resistance in pest species that metabolize unrelated chemical toxins.

In the current study, our overarching goal was to investigate whether fungicide exposure affects the phenotypic response of *L. decemlineata* to insecticides. We hypothesize that fungicides have a negative impact on *L. decemlineata* larval fitness, fungicides and insecticides can induce similar non-specific molecular detoxification mechanisms, and prior exposure to fungicides will impact the phenotypic response of *L. decemlineata* to insecticides. Herein, we investigated the acute and chronic fitness effects of two commonly used fungicides (chlorothalonil and boscalid) on *L. decemlineata* larvae. Building off these findings, we determined that prior-exposure to these fungicides can lead to a change in the phenotypic response in the insect when subsequently exposed to the insecticide imidacloprid. Finally, we investigated whether both fungicides and insecticides could induce similar mechanisms of metabolic detoxification. In this specific instance, we propose that a detoxification response to one chemical pressure (fungicide) could promote the detoxification of another chemical stressor (insecticide). This approach provides insight into the role select fungicides may play in development of insecticide resistance, and further provides a more biologically-based understanding of inadvertent selection events that affect insecticide (neonicotinoid) sensitivity. Our study suggests that the intensity of season-long disease management could have unequal impacts on *L. decemlineata* pesticide metabolism that, in turn, erodes the durability of key insecticides over time.

## Results

### Phenotypic Assay - Chronic and Acute Exposure to Fungicide

Larvae fed a continual diet of untreated *S. tuberosum* foliage developed normally and consistently gained weight over the 72 hour time course of the chronic exposure experiment. Larvae fed a continual diet of field-relevant rates of either chlorothalonil or boscalid did not consistently gain weight and comparatively lost weight at time points following exposure (p≤0.05), (**Fig. 1**). The assay was concluded after 72 hours because of the high mortality in both the chlorothalonil and boscalid groups (**Fig. S1**).

**Figure 1.**
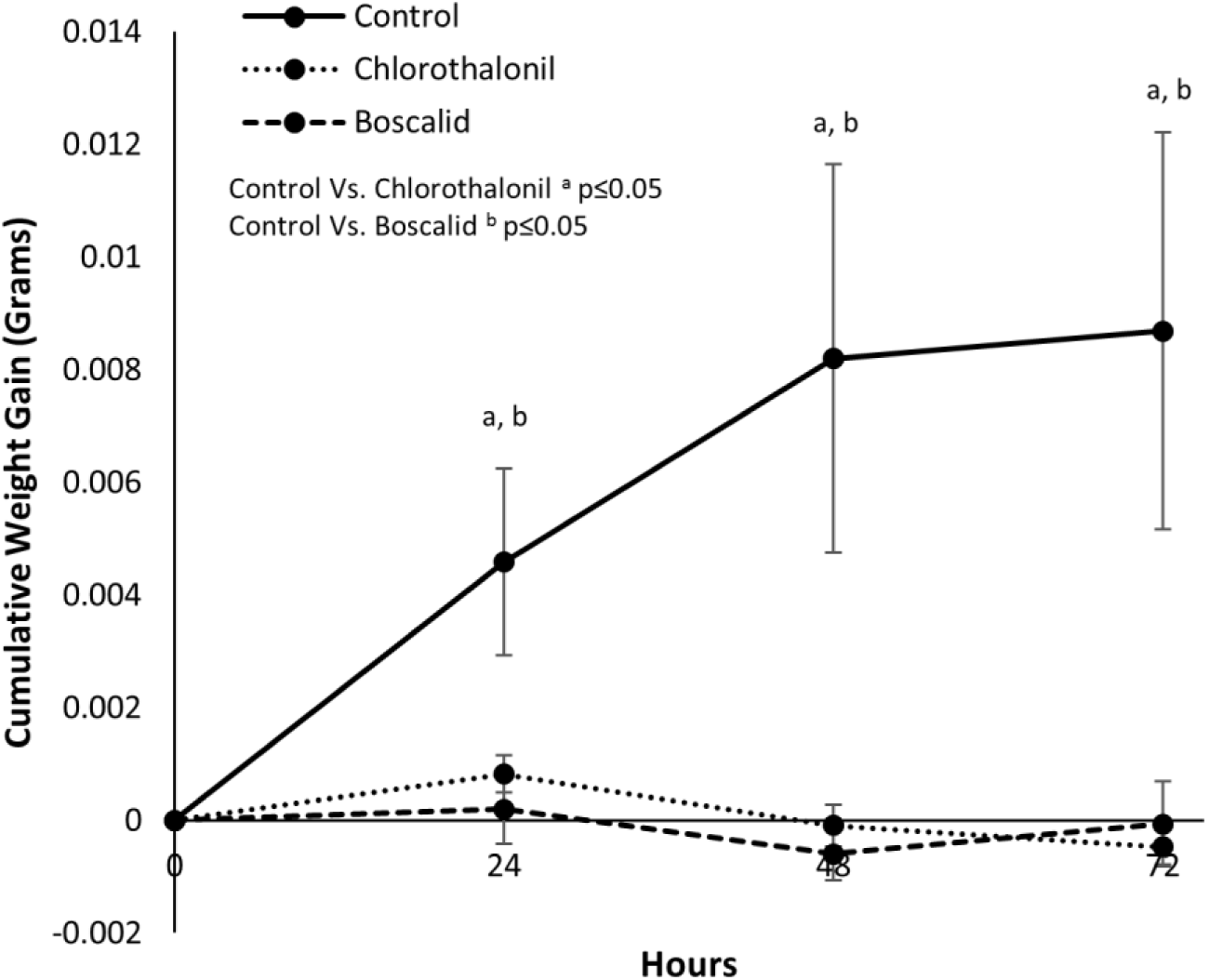
Effects of chronic exposure of chlorothalonil or boscalid on 2^nd^ instar larval weight gain over 72 hours (data represented as mean ± S.D.).

A one-time, acute exposure to chlorothalonil or boscalid did not impact weight gain in second instar larvae at most time-points measured (**Fig. 2**). However, weight gain after the topical application of chlorothalonil slightly, but significantly, decreased when compared to control at 24 (p=0.0008) and 144 (p=0.0237) hours after exposure. Weight gain after the topical application of boscalid slightly, but significantly, decreased 72 hours after exposure compared to the control (p=0.0165). Larval mass was tracked through the four instar molts as there was little mortality in all groups.

**Figure 2.**
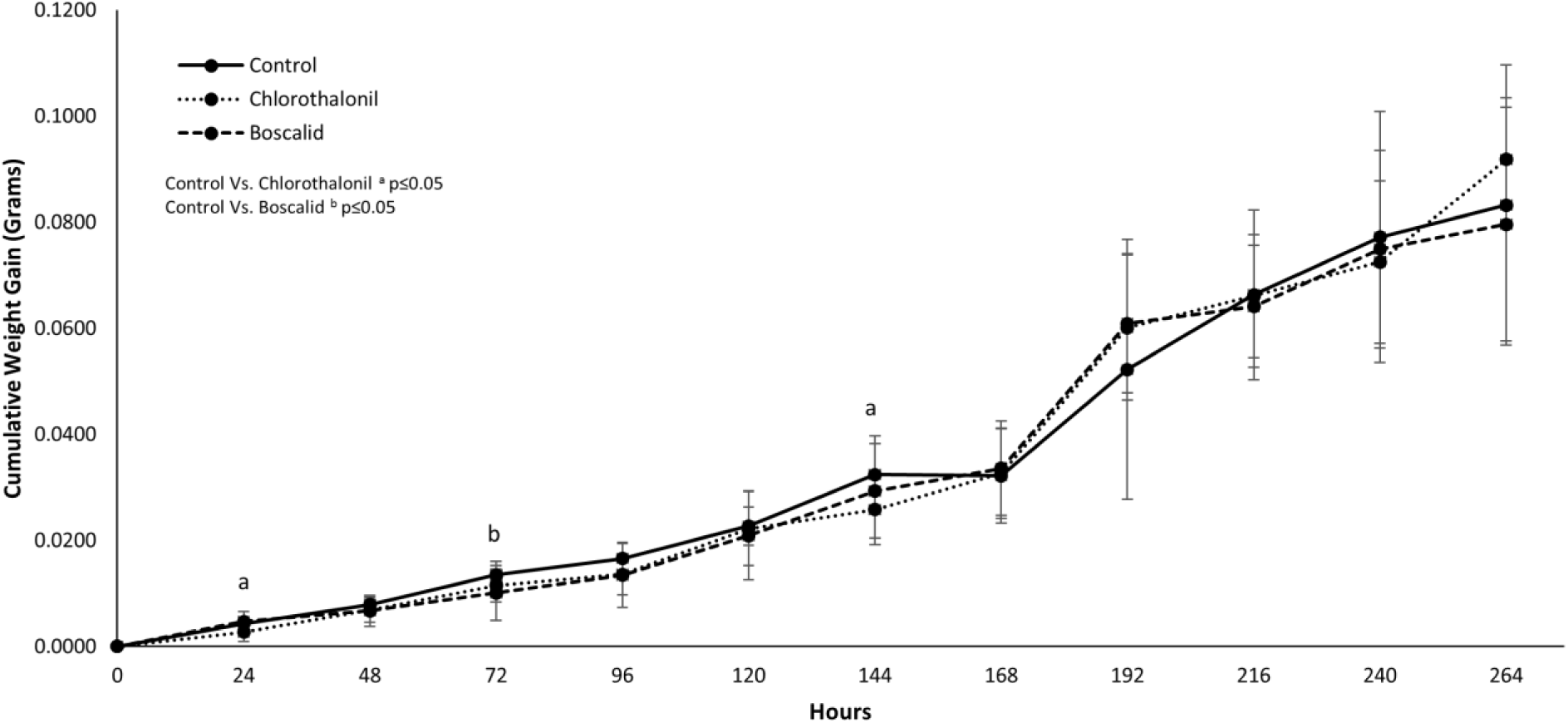
Effects of acute (one-time application) topical applications of chlorothalonil or boscalid on 2^nd^ instar larval weight gain over 264 hours (data represents mean ±S.D.).

### Imidacloprid Median Lethal Dose Assay

Methodology was successfully developed for a treated leaf disk feeding assay to assess the median lethal dose (LD_50_) of imidacloprid on the four larval and the adult life stages of *L. decemlineata*. Half lethal dose values for the 4 instar stages of beetles varied greatly. While there was statistical overlap between some life stages, trends in the data suggest that the first instar population was the most susceptible to imidacloprid while the adult stage was the least susceptible. LD_50_ values for each life stage, including the slope of the regression, 95% confident intervals, and chi-square values are presented in Table 1.

**Table 1.**
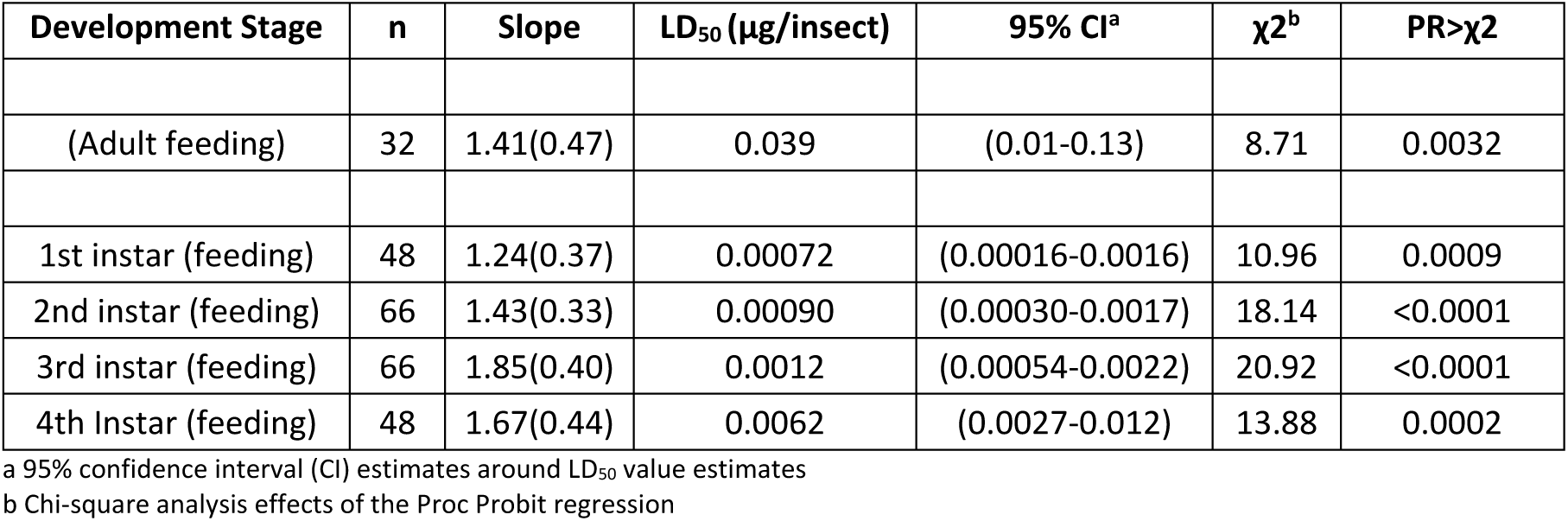
Estimated median lethal dose (LD_50_) estimates (expressed in µg per insect) for a laboratory-maintained, imidacloprid susceptible population (Arlington Agricultural Research Station, Arlington, Wisconsin) of *Leptinotarsa decemlineata* representing results from a feeding bioassay targeting the 4 larval instar stages and the adult stage.

### Phenotypic Response to Imidacloprid after Prior-Exposure to Fungicides

A field relevant, single application (prior-exposure) of either chlorothalonil or boscalid altered the phenotypic response of 2^nd^ instar *L. decemlineata* larvae to imidacloprid (Table 2). While the 95% confidence intervals overlap, trends in LD_50_ estimates suggest that prior-exposure to chlorothalonil decreased larval sensitivity to imidacloprid at both 2 and 6 hour time-points after topical application. In contrast, prior-exposure of *L. decemlineata* to boscalid comparatively increased imidacloprid sensitivity at similar time-points, resulting in lower estimated LD_50_ estimates for imidacloprid.

**Table 2:** Imidacloprid median lethal dose (LD_50_) estimates of *Leptinotarsa decemlineata* larvae previously exposed to either chlorothalonil or boscalid, and measured at 2 or 6 hour time-points following fungicide applications.

| Experimental Fungicide Treatment | n | Time Post-fungicide treatment (hours) | Slope | LD<sub>50</sub>(µg/insect) | 95% CI<sup>a</sup> | χ<sup>2</sup><sup>b</sup> | PR>χ<sup>2</sup> |
| --- | --- | --- | --- | --- | --- | --- | --- |
| Untreated control | 66 | NA | 1.43(0.33) | 0.00090 | (0.00030-0.0017) | 18.14 | <0.0001 |
| Topical application of chlorothalonil (6.9 µg/µl) | 66 | 2 | 1.83(0.41) | 0.0011 | (0.00050-0.0019) | 19.95 | <0.0001 |
| Topical application of chlorothalonil (6.9 µg/µl) | 66 | 6 | 2.80(0.64) | 0.0019 | (0.0010-0.0029) | 18.9 | <0.0001 |
| Topical application of boscalid (13 µg/µl) | 66 | 2 | 1.56(0.38) | 0.00030 | (0.000071-0.00067) | 16.4 | <0.0001 |
| Topical application of boscalid (13 µg/µl) | 66 | 6 | 1.02(0.28) | 0.00038 | (0.000039-0.00099) | 13.08 | 0.0003 |

a 95% confidence interval (CI) estimates around LD<sub>50</sub> estimates
b Chi-square analysis effects of the Proc Probit regression

### Glutathione S-Transferase Assay and Differential Transcript Abundance Analysis

Boscalid, chlorothalonil, and imidacloprid significantly (p≤0.05) increased glutathione S-transferase activity within *L. decemlineata* when compared to individuals fed untreated foliage. No glutathione S-transferase activity was detected in individuals fed untreated foliage. Levels of glutathione S-transferase induction rose significantly among groups of *L. decemlineata* exposed to either boscalid, chlorothalonil or imidacloprid (**Fig. 3**). Specifically, boscalid and imidacloprid induced significantly higher glutathione S-transferase activity in *L. decemlineata* when compared to chlorothalonil (p=0.0034 and p=0.0004 respectively) or control (p=0.0001 and p<0.0001 respectively), but were not significantly different from one another (p=0.2721). Chlorothalonil also induced significantly higher glutathione S-transferase activity in *L. decemlineata* when compared to control (p=0.0401).

**Figure 3:**
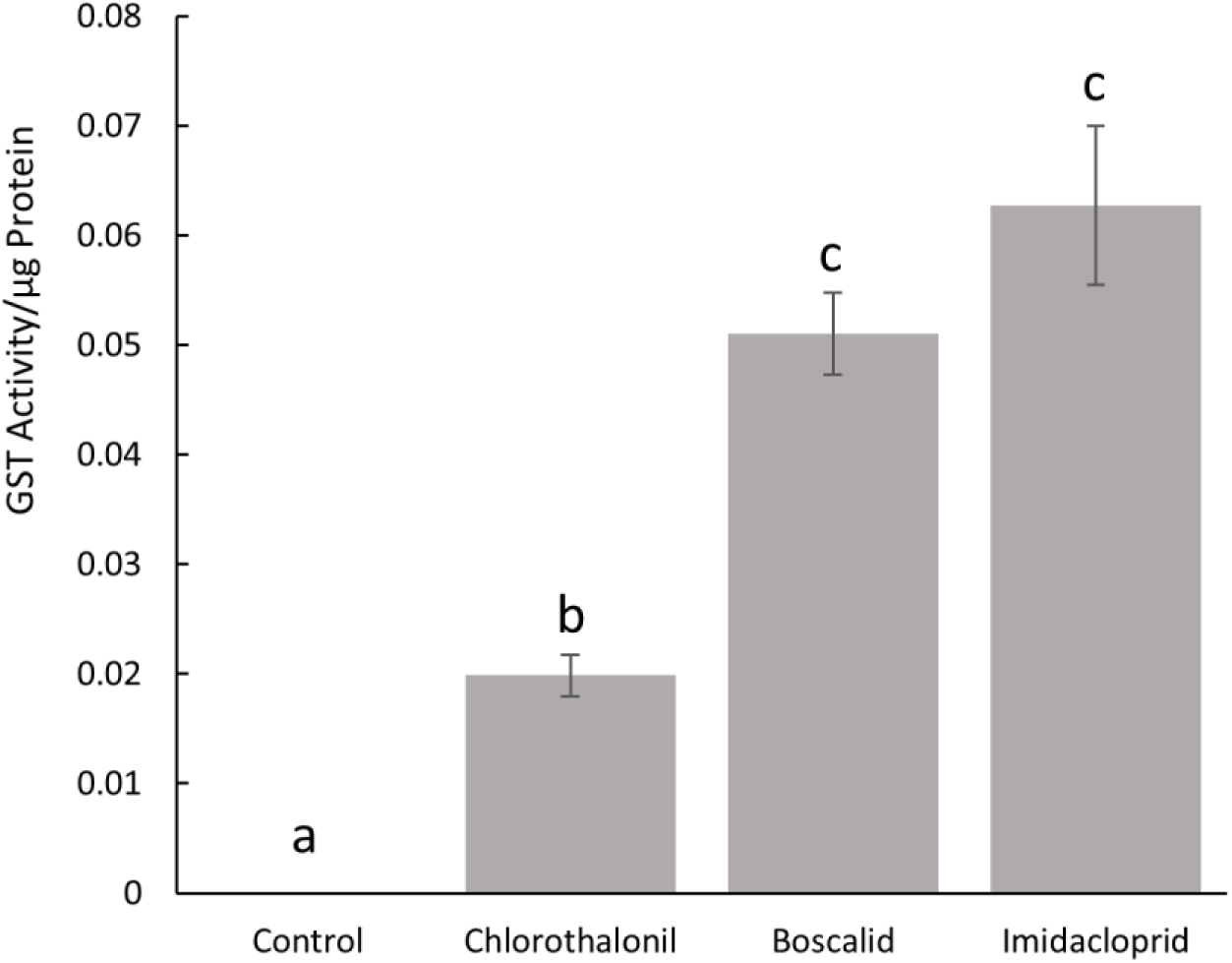
Levels of glutathione S-transferase induction (GST activity/µg protein) in *Leptinotarsa decemlineata* resulting from exposure to either chlorothalonil, boscalid, or imidacloprid (different letters indicate p≤0.05).

All three xenobiotic pesticides, chlorothalonil, boscalid, and imidacloprid, induced a significant fold change of a glutathione synthetase transcript, and both chlorothalonil and boscalid reduced the transcript abundance of a cytochrome p450 transcript (Table 3). The abundance of six transcripts was examined between a control group and those which received a prior exposure to either chlorothalonil, boscalid, or imidacloprid in order to determine if these xenobiotic exposures would specifically induce similar transcripts. A difference in fold change that was greater than two was considered significantly up or down regulated.

**Table 3:** Transcript abundance and associated fold change estimates in *Leptinotarsa decemlineata* resulting of exposure to either chlorothalonil, boscalid, or imidacloprid. Estimates were determined by transcript abundance analysis through quantitative PCR.

|  | Cuticular protein<br>(003961) | Cytochrome p450<br>(016769) | Cytochrome p450<br>(103658) | Cytochrome p450<br>(111691) | Glutathione synthetase<br>(114026) | Cytochrome p450<br>(115309) |
| --- | --- | --- | --- | --- | --- | --- |
| Control | 1 | 1 | 1 | 1 | 1 | 1 |
| Chlorothalonil | 1.09 | 1.01 | -1.28 | -1.29 | **2.72** | **-2** |
| Boscalid | -1.43 | -1.61 | -1.42 | 1.55 | **2.09** | **-3.41** |
| Imidacloprid | -1.47 | 1.45 | 1.05 | -1.17 | **3.66** | -1.59 |

Bolden values indicate significant positive or negative fold change $\geq 2$

### Pesticide Use Data

Complied data from USDA National Agricultural Chemical Use Survey from 1994-2014 ^15^ demonstrates trends in pesticide use throughout the United States. While the amount of neonicotinoid insecticides remains relatively consistent throughout agricultural potato producing regions, the amount of fungicides (chlorothalonil and boscalid) are higher in eastern and midwestern potato production regions when compared to the western potato production regions (**Fig 4**).

**Figure 4.**
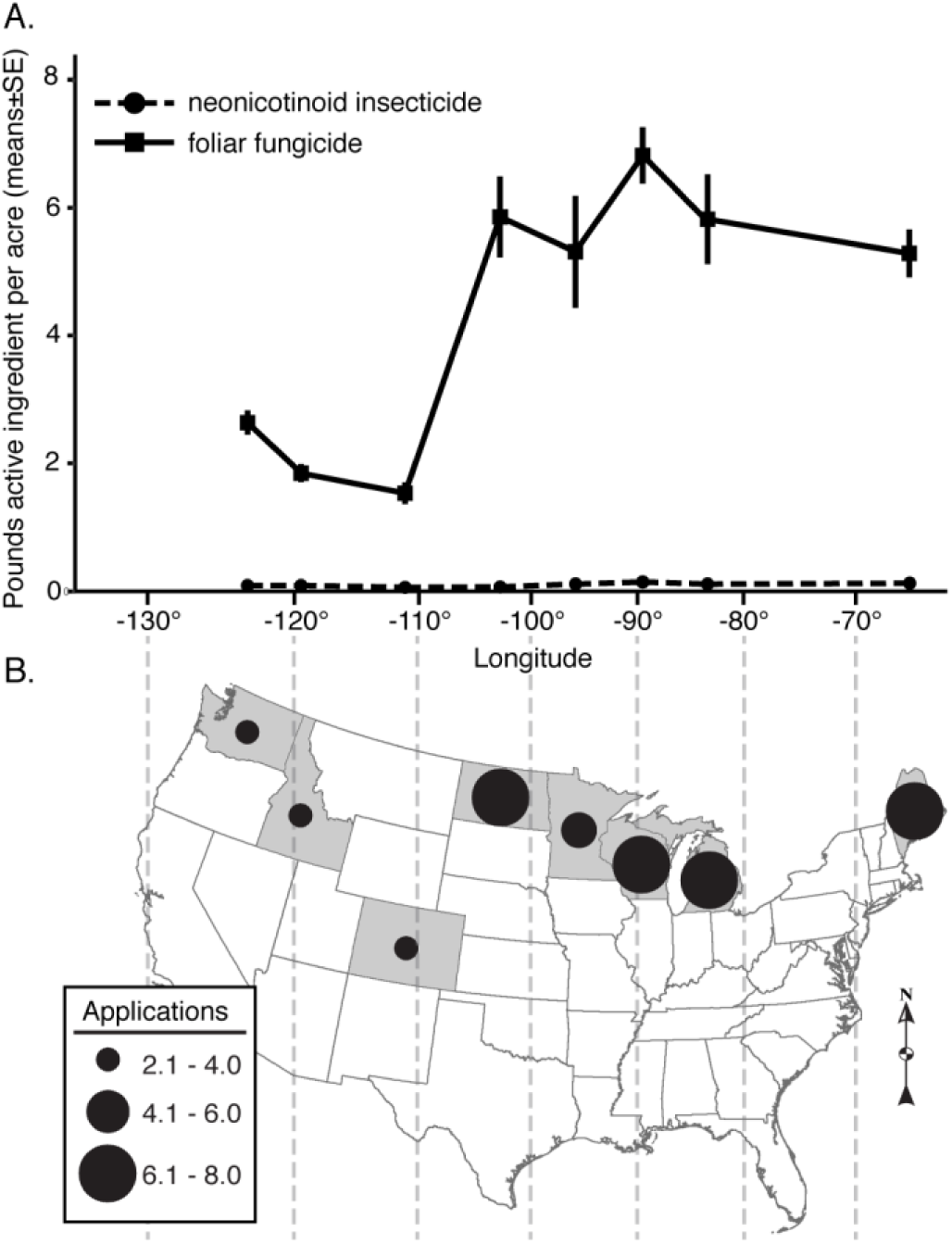
Average pounds active ingredient per acre of foliar fungicides and neonicotinoid insecticides applied to cultivated potato 1994-2014 (A). Average number of foliar fungicide applications during potato growing season (B) across different potato growing states. Values represent potato-specific pesticide amounts and application number reported from NASS 2014 ^15^.

## Discussion

The propensity of fungicides to induce the activation of molecular detoxification mechanisms in *L. decemlineata* is not well understood. Further, the ability of these fungicides to activate the same detoxification mechanisms that act on commonly used insecticides has not been well studied in the *L. decemlineata*. Here we demonstrate that a select set of common fungicides which are used to control both the early blight (*Alternaria solani*) and the late blight (*Phytophthora infestans*) pathogens, chlorothalonil and boscalid, have a significant fitness cost on *L. decemlineata* larvae, activate non-specific detoxification mechanisms, and influence the phenotypic response to the insecticide imidacloprid. The rapid turnover in insecticide registrations required to keep pace with the prevalence of insecticide resistance in *L. decemlineata* has direct impacts on the overall environmental footprint of potato pest management. Multiple insecticidal interventions are now commonplace in potato production regions where *L. decemlineata* resistance is a recurring issue and can be described as combinations of multiple, foliar and systemic insecticide treatments. In an effort to avert rapid resistance evolution, pest management practitioners have advocated for the rotation of chemical groups across sequential *L. decemlineata* generations and further suggest that the interval of re-exposure to specific MoA classes should be maximized to reduce the likelihood of resistance development ^16^. Multi-site mode of action fungicides, however, are often not rotated as frequently and are applied multiple times throughout each growing season ^15^. Without an understanding of the impacts of these fungicides on populations of *L. decemlineata*, a producer could be inadvertently hastening insecticide resistance even while attempting the use of best management practices for foliar pathogens.

The multi-site contact fungicide Chlorothalonil, first registered in 1966, has been a mainstay of fungicidal protection ^17,18^. Unfortunately, insect taxa in the agricultural field have also been exposed to chlorothalonil since 1966, allowing for 51 years of exposure to this xenobiotic to induce and activate many of the same genetic mechanisms that *L. decemlineata* use to detoxify insecticides. Boscalid, a carboxamide, with a targeted MoA of inhibiting succinate-dehydrogenase, was introduced more recently in 2003 ^18,19^ and has similar potential to inadvertently interact with insect populations. This compound has also been repetitively sprayed at high, labeled rates throughout potato production regions. The risk for fungal, bacterial and oomycete infections and their associated diseases in potato crops are the principal reasons for the prophylactic application of multiple fungicidal chemistries throughout the growing season. Although disease forecasting tools are regularly consulted to determine the risk for the occurrence or onset of these pathogens, once these established thresholds have been surpassed, serial applications of protective fungicides usually ensue. Moreover, as annual precipitation, humidity levels, and leaf wetness estimates are higher in Eastern and Midwestern potato production regions, the number of fungicidal applications increases sharply compared to western production regions. Concomitantly, there is considerable variation in the occurrence of insecticide resistance within and among populations of *L. decemlineata* across similar geographic regions of the US ^6,20^. Most notably, it has been well documented that populations of this pest species in Eastern and Midwestern potato production regions possess higher levels of measured insensitivity (resistance) to several insecticides when compared to populations in Western production regions ^5^. Specifically, sampled *L. decemlineata* populations from Idaho were 37X more susceptible to imidacloprid than populations collected in Maine^5^.

Figure 4 illustrates a more direct comparison of differences in the relative amounts of active ingredient applied and the frequency of application of fungicides and insecticides for eight major potato producing states from 1994-2014 ^15^. In Eastern (ME) and Midwestern (MI, WI, MN, and ND) potato production regions, it is common to have 5-7 foliar applications of fungicides in a single season, while the Northwestern region (WA, ID, and CO), has far fewer foliar applications (**Fig. 4B**). During seasons with high disease potential, Eastern and Midwestern crops may receive up to 10-12 successive, fungicide applications ^21,22^. While there is a clear trend in increased applications of fungicides in Midwest and Eastern regions when compared to the arid West, the overall amount of neonicotinoid insecticide applied to potato over this same interval remains relatively constant (**Fig. 4A**).

We hypothesize that greater levels of insecticide resistance to commonly used insecticides in the Midwest and Eastern potato production regions of the United States may be a consequence of higher and concomitant fungicidal inputs in these regions. Using a naive field-collected population of *L. decemlineata*, we demonstrated that fungicides have a significant impact on the fitness of this insect taxa and have the potential to induce multiple molecular detoxification mechanisms which are also utilized by *L. decemlineata* to combat insecticide treatments. By documenting that widely-used fungicides can induce similar detoxification mechanisms, we may be in a better position to modify best management practices to limit pesticide resistance in multiple insect taxa, including *L. decemlineata*.

From these investigations, we demonstrate the effects of acute (one-time application) and chronic (continuous) fungicide exposure at field relevant rates on a susceptible population of *L. decemlineata.* Throughout the current study, we focused on larval stages of *L. decemlineata*, as they are the target for many insecticides and are arguably the main life stage responsible for direct defoliation and plant damage. The design of the study was to mimic two possible field relevant scenarios. As fungicides are typically applied via foliar application, immature insects will necessarily be exposed to these residues on foliage. As a result, we investigated the effects of an acute, one-time topical application of fungicide. Here we observed that a one-time, acute dose had little observable effect on larval development. We next tested the effect of chronic exposure, where application would be evenly distributed throughout the field and growing season over successive time points. In our investigations, the chronic exposures have the potential to significantly influence larval fitness as evidenced by strong reductions in larval weight gain and arrested development. These observations suggest that larvae would have to adapt to survive these chronic and repetitive fungicidal inputs, as field populations are prolific in Midwestern and Eastern potato producing regions where these, and other fungicides, are regularly applied.

To determine whether prior-exposure of fungicides can change the phenotypic response of *L. decemlineata* to insecticides, we developed and subsequently implemented a novel, median lethal dose (LD_50_) feeding assay. Specifically, multiple doses of imidacloprid were topically applied to leaf disks using acetone as a carrier solution and all of the acetone was allowed to evaporate prior to infestation. The area of the leaf disk (e.g. 0.13 cm^2^) used for these investigations was designed to be less than the average consumption rates typical for an early instar larva over a 24 hour period. In this way, test insects consistently consumed the entire leaf disk over the 24 hour exposure interval. For the lower, sub-lethal doses, this was always the case, but as larvae were exposed to increasingly higher doses, mortality often occurred before complete consumption of the leaf disk. From these observations, we still concluded that the doses would have been lethal if the entire disk had been consumed. Using different doses of imidacloprid, we accurately determined the LD_50_ values for the naïve field strain of *L. decemlineata* population used in these investigations (Arlington Agricultural Research Station population, Arlington, WI), and across all larval instar stages and the adult stage. Even though there was a slight overlap in 95% confident intervals, we demonstrated that the amount of insecticide required to result in median mortality varied among different life stages. From these results, we chose to focus on the 2^nd^ instar larval stage to further examine whether prior-exposure to chlorothalonil or boscalid would influence the phenotypic response of early larvae to imidacloprid as measured by LD_50_ estimates at both 2 and 6 hours after exposure to the fungicide. A relatively short time course was chosen to mimic field relevant topical exposure to fungicides, and corresponding feeding on insecticide treated foliage. Prior-exposure to the fungicide resulted in unique trends in phenotypic response to imidacloprid. A prior-exposure to boscalid markedly decreased the amount of imidacloprid required to kill half the test population, whereas prior-exposure to chlorothalonil resulted in an elevated level of insecticide required to kill half of the test population. While again there was a slight overlap in 95% confident intervals, trends in these data suggest that chlorothalonil and boscalid can differentially influence the relative fitness of *L. decemlineata* populations in advance of insecticide exposures. Further studies are needed to observe whether long-term chronic exposure would have similar phenotypic results.

The activation of nonspecific enzymes, which attack functional groups rather than the specific molecules, are induced by both chlorothalonil and boscalid. One such nonspecific enzyme is a phase 2 conjugating protein known as glutathione S-transferase. Glutathione S-transferase can catalyze the conjugation of glutathione to a xenobiotic compound rendering it more easily excreted, and in turn, less toxic ^23^. We detected significantly elevated levels of glutathione S-transferase induced by chlorothalonil, boscalid and imidacloprid compared to a control (**Fig. 3**). The upregulation of a specific enzyme which metabolizes pesticides indicates that fungicides may play a role in increasing pesticide resistance in *L. decemlineata.* We identified 6 other potential genetic targets that have been suggested to play a role in insecticide resistance in *L. decemlineata* ^8,9^, a cuticular protein, a glutathione synthetase, and four cytochrome p450s. While the targets that were chosen are only a small fraction of those which encode potential mechanisms of detoxification, many of the targets have been previously shown to be constitutively upregulated in insecticide resistant populations of *L. decemlineata* in Wisconsin ^8,9^. After exposure to chlorothalonil, boscalid, or imidacloprid, we observed significant trends in transcript regulation in two of the genetic targets. The first was the glutathione synthetase, which is a vital enzyme in glutathione production. Chlorothalonil, boscalid, and imidacloprid all upregulated the transcripts corresponding to this enzyme, suggesting that select fungicides and the insecticide imidacloprid are inducing this enzyme to increase glutathione for glutathione S-transferase metabolism. The second was the downregulation of transcript (115309) corresponding to a cytochrome p450 9Z26. Both boscalid and chlorothalonil significantly down regulated theses transcripts, and imidacloprid also trended towards down regulation. It is commonly known that insects have cost-benefit tradeoffs in enzymatic activation, and enzymatic detoxification requires energy expenditure, and while certain transcripts are upregulated, others will be down regulated as a response ^24^. Importantly, fungicides and imidacloprid are influencing transcript abundance after exposure in a similar manner.

This study demonstrates the phenotypic effects that chlorothalonil and boscalid have on *L. decemlineata* 2^nd^ instar larvae, including the effect of prior-exposure to fungicides that led to changes in the relative toxicity of the insecticide imidacloprid. We have further demonstrated the induction of similar non-specific detoxification mechanisms between fungicides and insecticides through enzymatic assays and transcript abundance studies. Although these results support our hypothesis that prior exposure to fungicides may be influencing interactions between *L. decemlineata* and insecticides, more work is required to better understand the specific mechanisms. RNA sequencing experiments examining the overall transcript abundance induced by similar xenobiotic pesticides will be vital to examine these important genetic responses. Furthermore, other environmental and human inputs, such as herbicides, insecticides, and fertilizers, may also drive insecticide resistance to varying degrees, and need to be taken into account when inferring the relative contributions of these investigations in the evolution of insecticide resistance. The data presented here were obtained from larvae representing a susceptible field population maintained as laboratory strain. The naïve, susceptible strain provides a baseline for future investigations, however, future studies could extrapolate more data by expanding similar methodology onto an insecticide resistant field population and confirming transcript abundance for similar genetic mechanisms.

As growers and pest managers face the risk of increasingly resistant insect pest populations, they need to understand the ramifications of the chemical inputs commonly used in agricultural fields. Our study demonstrates that select xenobiotic inputs have an effect outside of their targeted species. Furthermore, the fact that the fungicidal programs have been implemented for more than 50 years in select areas of the US suggest that any arthropods exposed to these chemistries have had substantial time to adapt. We suggest that some fungicides may be partially contributing to measured levels of insecticide resistance in *L. decemlineata* by demonstrating that fungicides have a significant fitness cost on *L. decemlineata* larvae, activate non-specific detoxification mechanisms, and influence the phenotypic response to the insecticide imidacloprid. Furthermore, growers should consider the impact of these multi-site mode of action fungicides in advance of insecticidal applications, as general molecular detoxification mechanisms will likely be activated from the fungicide, rendering the insecticides less effective in future applications.

## Materials and Methods

### Housing and Maintenance of Lab Colony

Approximately 300 adult beetles were collected on June 20^th^ 2016 from the Arlington Agricultural Research Station, Arlington, Wisconsin (AARS, 43.315527, −89.334545), where populations have little exposure to insecticides. Beetles in this population remain highly susceptible to imidacloprid ^8,16^. Healthy adult beetles were hand-collected from the canopy of potato plants, placed in plastic containers and returned to the University of Wisconsin-Madison. Beetles were sustained on healthy potato plants in mesh cages under a 16:8 hour light:dark (L:D) photoperiod. Untreated foliage from potato plants was obtained from plants grown at the University of Wisconsin-Madison Greenhouse and provided to beetles daily. Adult beetles were given the opportunity to randomly mate and lay clutches of eggs on potato foliage. Egg masses were collected daily and placed on filter paper in 100 mm petri dishes (Corning, Corning, New York) and held at 26°C, 70% relative humidity (RH), and 16:8 (L:D) photoperiod. Following egg hatch, larvae were provided untreated foliage daily and maintained as cohort groups through the remainder of their larval development before being returned to mesh cages with fresh potato plants and soil and allowed to pupate and subsequently emerge as adults.

### Phenotypic Assay - Chronic and Acute Exposure to Fungicide

From the previously described lab colony, 120, second instar larvae were identified according to Boiteau et al. ^25^. Field-relevant levels of technical chlorothalonil (Syngenta Crop Protection, Basel, Switzerland) and boscalid (Tokyo Chemical Industry, Chennai, India) were dissolved in acetone at concentrations of 6.9 µg/µl and 13 µg/µl, respectively, which also corresponded to application rates of 206 g ha^-1^ of chlorothalonil and 707 g ha^-1^ of boscalid. Individual larvae were placed in a single well of a flat-bottom, 12 well Falcon plate (Corning Inc., Corning, New York). Each well contained a water-dampened sponge, covered by filter paper giving the larva a platform on which to stand and feed. For the chronic exposure assay, potato foliage leaf disks (2.01 cm^2^) were dipped in the field relevant solutions of boscalid or chlorothalonil in acetone, allowed to completely dry, and were then presented to the second instar larvae. Every 24 hours the remnants of the previous leaf disk were removed and a newly treated leaf disk was presented to each larva (n=20 larvae per treatment), and the experiment was conducted for 72 hours. For the acute exposure assay, second instar larvae were topically dosed only once with field relevant solutions described above of chlorothalonil, boscalid, or an acetone control (n=20 per treatment). Specifically, 1 μL of solution was topically applied to the dorsal surface of the larva’s abdomen and allowed to completely absorb into the insect cuticle (n=20 larvae per treatment). After topical application, individual larvae were placed onto untreated potato leaf disks in a single well of a flat-bottom, 12 well, Falcon plate, as described previously. Every 24 hours the remnants of the previous leaf disk were removed and a new, untreated leaf disk was provided to each larva. Insects in their experimental plates were held at 26°C, 70% RH, and a 16:8 (L:D) photoperiod until the experiment was concluded. To track the progress of larval development, each larva was weighed using an AE 100 analytical balance (Mettler Toledo, Columbus, OH) every 24 hours through all larval development stages and a one-way ANOVA with a Tukey, post-hoc analysis was performed to determine significant changes in weight gain, with p≤0.05 considered significant.

### Imidacloprid Median Lethal Dose Assay

Feeding bioassays were developed to generate estimates of the median lethal dose (LD_50_) to imidacloprid. In 12 well, Falcon microplates, replicate sets of 0.13 cm^2^ leaf disks were placed upon a damp sponge covered by a piece of filter paper. The sponge and filter paper were arranged to take up approximately ½ of the volume of each well and were used to maintain a high level of humidity. A single *L. decemlineata* was placed into each well. Immediately prior to placement of the insect, 1 µl aliquots of technical grade imidacloprid (Bayer Crop Science, Kansas City, MO) in acetone ranging between 0 to 0.39 µg imidacloprid were applied to leaf disks (n=4-6 per dose). After application to the leaf disk, the acetone was allowed to evaporate leaving only the insecticide residue before the insect was placed in the well. Each insect was given the opportunity to consume the leaf disk over a 24 hour period and percent mortality was observed. Assay plates were held at 26°C, 70% RH, and a 16:8 (L:D) photoperiod.

Adult *L. decemlineata* mortality was determined by presenting the beetle the opportunity to climb a pencil: if they could move a full body length they were considered alive, if they appeared alive, but could not move a body length then they were considered incapacitated, and if they had no movement, even after pinching metathoracic legs with tweezers, they were considered dead ^26^. To determine larval mortality, each larva was inverted onto its dorsal side. Unaffected larvae could invert themselves within 20 seconds, incapacitated larvae could not invert themselves within 20 seconds and dead larvae showed no movement. Incapacitated and dead individuals were pooled and LD_50_ values were calculated using probit regression analysis (PROC PROBIT, SAS Institutes).

### Phenotypic Response to Imidacloprid after Prior-Exposure to Fungicides

To determine if prior-exposure to fungicides could influence the phenotypic response to imidacloprid, a total of 264, second instar larvae were initially sorted. Upon initiation of the experiment (t=0), 132, second instar larvae were topically dosed with 1 µl of a field relevant concentration of either chlorothalonil (6.9 µg/µl in acetone) or boscalid (13 µg/µl in acetone) (132 individuals per fungicide). Specifically, 1 μL of solution was topically applied to the dorsal surface of the larva’s abdomen and allowed to completely absorb through the cuticle, as described previously. After topical application, larvae were placed in an incubator at 26°C, 70% RH for 2 and 6 hour periods. At each of these corresponding time points, 66 larvae were removed representing each fungicide treatment and an imidacloprid LD_50_ feeding bioassay was subsequently conducted as previously described. Median lethal dose estimates were calculated using probit regression analysis (PROC PROBIT, SAS Institutes).

### Glutathione S-Transferase Assay and Differential Transcript Abundance Analysis

At the initiation of this experiment (t=0), 96, 2^nd^ instar larvae were sorted and initially starved for 4 hours. During this period, larvae were placed at 26°C and 70% RH. After the 4 hour starvation period, larvae were broken into 4 groups (n=24/group). Individual larvae were placed in a single well of a 12-well, flat-bottom plate containing a leaf disk (0.13 cm^2^) with either a dried acetone control, 13 µg of boscalid, 6.9 µg of chlorothalonil, or 0.000078 µg imidacloprid. Larvae were fed treated leaf disks over a 24 hour time period, after which time they were sorted into 6 biological replicates of 4 pooled individuals representing each experimental group. Three replicates (n=4 pooled individuals/replicate) for each group were later used for a glutathione S-transferase activity assay while the remaining 3 replicates (n=4 pooled individuals/replicate) for each group were used for transcript abundance analysis. To determine glutathione S-transferase activity, whole larvae were sacrificed and tissue was homogenized according to Cayman Chemicals, Glutathione S-Transferase assay kits instructions (Cayman Chemicals, Ann Arbor, MI). A bicinchoninic acid assay (BCA assay) (ThermoFisher Scientific, Waltham, MA) was run to standardize protein concentrations between the tissue homogenates. A Glutathione S-Transferase assay was then conducted according to the kit guidelines (Cayman Chemicals, Ann Arbor, MI). Readings below the detection limit were recorded as zero. A one-way ANOVA with a Tukey post-hoc analysis was run to determine significant changes in activity, with p≤0.05 considered significant. Transcript abundance of six target genes previously shown to be up-regulated in imidacloprid-resistant populations of *L. decemlineata* were examined to observe if prior exposure to fungicides could also induce similar transcripts ^8,9,27^. Total RNA was extracted from each larval group with Trizol (Life Technology, Grand Island, NY). DNA contamination was removed with TurboDNase (Life Technology, Grand Island, NY, USA) and total RNA was purified through EtOH precipitation, air dried until no visible liquid was observed, and then suspended in 50 µL DNase/RNase-free H_2_O. All RNA concentrations were equalized before input into the cDNA synthesis kit, and the subsequent cDNA was generated with a Super Script III kit (ThermoFisher Scientific). The cDNA was diluted to a final concentration of 5 ng/µL RNA equivalent input for qPCR. Rp4 was used as a reference gene in the analysis ^28^. The qPCR reaction was run on a CFX-96 platform (Bio-Rad Laboratories, Hercules, CA, USA) with a master mix of Bullseye EverGreen (MIDSCI, Valley Park, MO, USA). The qPCR reactions were conducted using the Pfaffl efficiency calibrated methodology; primer and primer efficiency (amplification efficiency of reactions as described by Pfaffl ^29^) are found in the supplementary information (Table S1). Triplicate reactions were run at 95 °C for 10 min, followed by 95 °C for 15 s, and 62 °C for 60 s for a total of 40 cycles. Mean Ct values (Table S2) were collected for each biological replicate and fold change estimates between the non-xenobiotic exposure and the three pesticides were calculated.

### Pesticide Use Data

To determine the relationship between fungicide and insecticide use in US potato, we compiled publicly available potato-specific pesticide use data reported in the USDA National Agricultural Chemical Use Survey from 1994-2014 ^15^. For this period, we first analyzed application reports for three neonicotinoids targeting *L. decemlineata* (clothianidin, imidacloprid, thiamethoxam) and two common foliar fungicides studied here (boscalid, chlorothalonil). To determine the frequency of potential fungicide exposure, we summarized the average number of applications of boscalid and chlorothalonil for major producing states over the 21 y period. We also calculated the average pounds active ingredient per hectare for these potato fungicides and insecticides. Annual estimates for application number and amount sprayed were summed across active ingredients within both pesticide group. The average number of applications and average pounds of active ingredient per acre (±SE) was calculated for each state using R (R-Core Development Team, Version 3.2.3). The average number of fungicide applications was mapped to state centroids using ArcGIS (Version 10.1, ESRI, Redlands, CA USA).

## Compliance with ethical standards

### Ethical approval

This article does not contain studies with any human participants and no specific permits were required for field collection or experimental treatment of *L. decemlineata* for the study described.

## Acknowledgments

The authors would like to acknowledge support from Wisconsin Potato and Vegetable Growers Association and the associated agricultural community. This research was partially supported by a combination of funding from: (1) USDA-NIFA-ARFI-ELI Postdoctoral Fellowship Grant 12105364 (2) University of Wisconsin, Hatch Formula 142 Funds (142PRJ72SV).

