## Supplementary Materials for "Agricultural fungicides inadvertently influence the fitness of Colorado potato beetles, *Leptinotarsa decemlineata*, and their susceptibility to insecticides"

Supplementary Table 1. Quantitative PCR primers and primer efficiency, transcript identification including associated BLASTx NCBI accession number referenced by Clements et al. [8,9,27].

| | Forward Primer (5'-3') | Reverse Primer (5'-3') | Primer Efficiency | Transcript BLAST x Result | NCBI Accession Numbers |
|---|---|---|---|---|---|
| RP4 (Reference) | AAAGAAACGAGCATTGCCCTTCCG | TTGTCGCTGACACTGTAGGGTTGA | 1.93 | | |
| LDEC003961 (Cuticular protein) | ACCTGCTGCCGGTATTATTG | TACAGTTCCAGAGGGTCCAG | 1.96 | Cuticular protein | XP_966639.1 |
| LDEC016769 (Cytochrome p450) | CAGGTCTGACAAGGATATGGTTAG | TCCAGAGCTTTCGGATGATTC | 1.98 | Cytochrome p450 | XP_973153.1 |
| comp103658 (Cytochrome p450) | TCCTCACTGAATCTTTCTGGATCG | AGCCCAGATGAGAAGCCATTAC | 2.02 | Cytochrome p450 9z4 | NP_001164248 |
| comp111691 (Cytochrome p450) | TGCCGTCTCCTAGCTTTGTAAC | CGGATTCGATACCAGTTCAACAC | 2.05 | Cytochrome p450 monooxygenase | XP_972348 |
| comp114026 (Glutathione synthetase) | CAGAGCAGGGTATGAACCTAATC | CCAGCCAAGTGATACTGAATCG | 1.97 | Glutathione synthetase | XP_968070 |
| comp115309 (Cytochrome P450) | CGAGAAATGCGACCTATTCTCAG | ACACAGTCTTGGTCTTTCTTGAG | 1.98 | Cytochrome P450 9Z26 | KJ476503.1 |

Supplementary Table 2. Mean CT values determined by quantitative PCR used to calculate transcript expression. Each transcript of interest was run on a separate CFX 96 plate with the corresponding reference transcript RP4 (to account for inter-assay variability).

| | **Control (leaf disk only)** | **Fungicide ( chlorothalonil + leaf disk)** | **Fungicide (boscalid + leaf disk)** | **Insecticide (imidacloprid + leaf disk)** |
|---|---|---|---|---|
| | **Mean CT ±SD** | **Mean CT± SD** | **Mean CT± SD** | **Mean CT±SD** |
| RP4 (Reference for LDEC003961) | 19.62±0.86 | 18.90±0.45 | 18.74±0.34 | 18.96±0.31 |
| LDEC003961 (Cuticular protein) | 33.95±2.81 | 33.12±0.93 | 33.63±1.53 | 33.89±2.26 |
| RP4 (Reference for LDEC016769) | 18.39±0.31 | 17.95±0.33 | 18.14±0.20 | 18.55±0.24 |
| LDEC016769 (Cytochrome p450) | 34.24±0.33 | 33.80±0.20 | 34.70±0.61 | 33.85±1.39 |
| RP4 (Reference for comp103658) | 18.67±0.41 | 18.16±0.38 | 18.41±0.38 | 18.57±0.18 |
| comp103658 (Cytochrome p450) | 21.66±0.15 | 21.54±0.26 | 21.92±0.60 | 21.50±0.17 |
| RP4 (Reference for comp111691) | 19.14±1.30 | 18.75±0.52 | 19.30±1.03 | 18.43±0.22 |
| comp111691 (Cytochrome p450) | 23.38±0.09 | 23.38±0.28 | 22.92±0.55 | 22.96±0.35 |
| RP4 (Reference for comp114026) | 18.53±0.41 | 17.95±0.31 | 18.20±0.36 | 18.39±0.18 |
| comp114026 (Glutathione synthetase) | 22.03±0.81 | 20.91±0.18 | 21.54±0.38 | 20.90±0.42 |
| RP4 (Reference for comp115309) | 18.71±0.38 | 18.19±0.32 | 18.39±0.20 | 18.76±0.26 |
| comp115309 (Cytochrome P450) | 18.64±0.68 | 19.16±0.30 | 20.13±0.73 | 19.37±0.41 |

Supplementary Figure 1. Percent survivorship of second instar larval exposed to chronic doses of field relative rates of chlorothalonil, boscalid and a no treatment control.

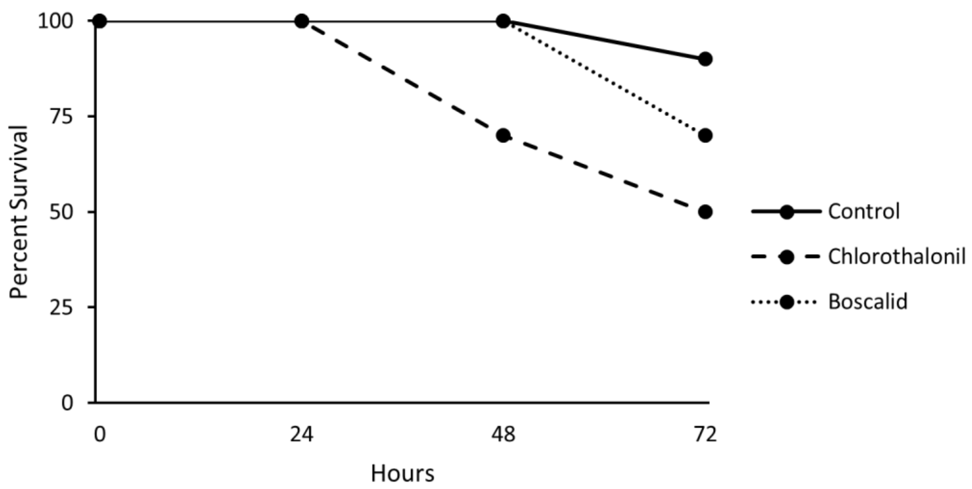
